# BiG-SLiCE: A Highly Scalable Tool Maps the Diversity of 1.2 Million Biosynthetic Gene Clusters

**DOI:** 10.1101/2020.08.17.240838

**Authors:** Satria A. Kautsar, Justin J. J. van der Hooft, Dick de Ridder, Marnix H. Medema

## Abstract

**Background:** Genome mining for Biosynthetic Gene Clusters (BGCs) has become an integral part of natural product discovery. The >200,000 microbial genomes now publicly available hold information on abundant novel chemistry. One way to navigate this vast genomic diversity is through comparative analysis of homologous BGCs, which allows identification of cross-species patterns that can be matched to the presence of metabolites or biological activities. However, current tools suffer from a bottleneck caused by the expensive network-based approach used to group these BGCs into Gene Cluster Families (GCFs).

**Results:** Here, we introduce BiG-SLiCE, a tool designed to cluster massive numbers of BGCs. By representing them in Euclidean space, BiG-SLiCE can group BGCs into GCFs in a non-pairwise, near-linear fashion. We used BiG-SLiCE to analyze 1,225,071 BGCs collected from 209,206 publicly available microbial genomes and metagenome-assembled genomes (MAGs) within ten days on a typical 36-cores CPU server. We demonstrate the utility of such analyses by reconstructing a global map of secondary metabolic diversity across taxonomy to identify uncharted biosynthetic potential. BiG-SLiCE also provides a "query mode" that can efficiently place newly sequenced BGCs into previously computed GCFs, plus a powerful output visualization engine that facilitates user-friendly data exploration.

**Conclusions:** BiG-SLiCE opens up new possibilities to accelerate natural product discovery and offers a first step towards constructing a global, searchable interconnected network of BGCs. As more genomes get sequenced from understudied taxa, more information can be mined to highlight their potentially novel chemistry. BiG-SLiCE is available via https://github.com/medema-group/bigslice.

## Background

The microbial world is teeming with diverse microorganisms competing and collaborating for survival. A major theme in these microbial interactions is the use of bioactive compounds from secondary metabolism. Some of these compounds have long been exploited by humans for their medicinal, antifungal, and antibacterial effects [1]. Some others found their use in agriculture [2], wastewater treatment [3], and everyday products such as detergents and cleaning products [4]. A recent report by the World Health Organization (WHO) highlights the need to explore novel chemistry from nature amid the increasing problems caused by antimicrobial-resistant (AMR) bacteria [5]. It was previously estimated that there might be billions of microbial species living on earth [6,7] and even from the heavily mined genus of *Streptomyces*, novel discoveries continue to be made [8–13]. Due to the sheer size of microbial and enzymological biodiversity, there exists a vast repertoire of potentially useful compounds remains to be unearthed. More fundamentally, by learning about microbes and the compounds they produce, we can gain knowledge about mechanisms of interaction within microbiomes, enabling us to study how their microbial composition is associated with human health and disease [14] or to learn about the symbiotic relationships between soil microbes and their plant host [15].

One promising way to reveal this knowledge is to leverage the power of large-scale omics. Metabolomics provides a complete snapshot of metabolites produced by microbes at a given time, while transcriptomics and proteomics provide insight into metabolic pathways and their regulation [16–18]. On the other hand, genomics allows the rapid profiling of an organism’s metabolic potential via the computational prediction of Biosynthetic Gene Clusters (BGCs) [19–21]. Previous studies [22–29] show that grouping BGCs with similar architecture (i.e. sharing a similar set of homologous core genes) into Gene Cluster Families (GCFs) can yield useful insights into the chemical diversity of the analyzed strains, and can support linking BGCs to their products via the emerging technique of metabologenomics [23,25]. BGCs responsible for the production of retimycin A [27], tambromycin [25], tyrobetaines [30] and several detoxin-rimosamide analogs [22] have been elucidated via this approach. GCFs have also been used as functional markers in human health studies [31,32] and to study soil suppressiveness against fungal pathogens [33]. This gradual shift from a gene-centric approach in functional metagenomics to a gene cluster-centric one is likely to be stimulated further with the increasing accessibility of long sequencing reads that easily span tens to hundreds of kbp (kilobase pairs) in size [34], effectively covering the full span of a typical microbial BGC within a single read.

Given their direct relationship to the catalytic enzymes, and subsequently, the compounds produced from their encoded pathways, BGCs (and, by extension, GCFs) can serve as a proxy to explore the chemical space of microbial secondary metabolism. By cataloging all the GCFs in sequenced microbial genomes, one can obtain an overview of the existing chemical diversity and gain insights into what future lead discovery efforts should prioritize. For example, one could focus on species harboring the most potential novelty, or on identifying natural variants of a known antibiotic-producing BGC. For such global analyses, the clustering algorithm to group BGCs into GCFs needs to be able to work with massive volumes of data. While a trend of increasing input capacity can be observed for the past 5 years (from 11,000-33,000 analyzed BGCs in 2014 [23,24] to 73,260 in 2019 [22]), it is still dwarfed by the total amount of data currently available. As of 27 March 2020, antiSMASH-DB [35] and IMG-ABC [36], the two largest BGC databases, jointly comprise 565,096 BGCs predicted from 85,221 bacterial genomes. This number will increase even more if we account for genomes and metagenomes not covered by these databases. For example, assuming they hold similar average numbers of BGCs, the ~180,000 bacterial genomes in the NCBI RefSeq database (https://www.ncbi.nlm.nih.gov/refseq/) may yield more than a million BGCs when processed with tools like antiSMASH.

To handle a dataset this large, even the currently fastest tool (one tool we previously developed, BiG-SCAPE [22]) will require an estimated 37,000 hours of runtime on a 36-core CPU (see Results and Discussion), which is impractical if not impossible. A major bottleneck is the expensive pairwise BGC comparison used to construct similarity networks and perform clustering analysis, leading to quadratic time complexity (*O(n*^2^), where *n* is the total number of BGCs). Thus, there is an urgent need for an alternative method that better scales with the available genomic data, which will grow even further as the cost and performance of Next Generation Sequencing (NGS) technology continue to improve and get democratized [37]. Here, we introduce BiG-SLiCE (*Biosynthetic Genes Super-Linear Clustering Engine*), which projects BGCs into Euclidean space to enable the usage of a partitional clustering algorithm running in a near-linear (~*O(n)*) time complexity. Using this approach facilitates analysing large datasets of BGCs orders of magnitude faster, finally allowing truly global GCF analyses on all available microbial genomes.

## Methods and Implementation

The BiG-SLiCE workflow starts at the vectorization (feature extraction) step [Figure 1A], converting input BGCs into vectors of numerical features based on the absence/presence and bitscores of hits obtained from querying BGC gene sequences against a library of curated profile Hidden Markov Models (pHMMs). Those features are then processed by a super-linear clustering algorithm [Figure 1B], resulting in a set of centroid feature vectors representing the GCF models. All BGCs in the dataset are finally queried back against those models [Figure 1C], outputting a list of GCF membership values for each BGC. In the end, an interactive visualization output is produced, which enables users to explore the analyzed data [Figure 1D].

**Figure 1.**
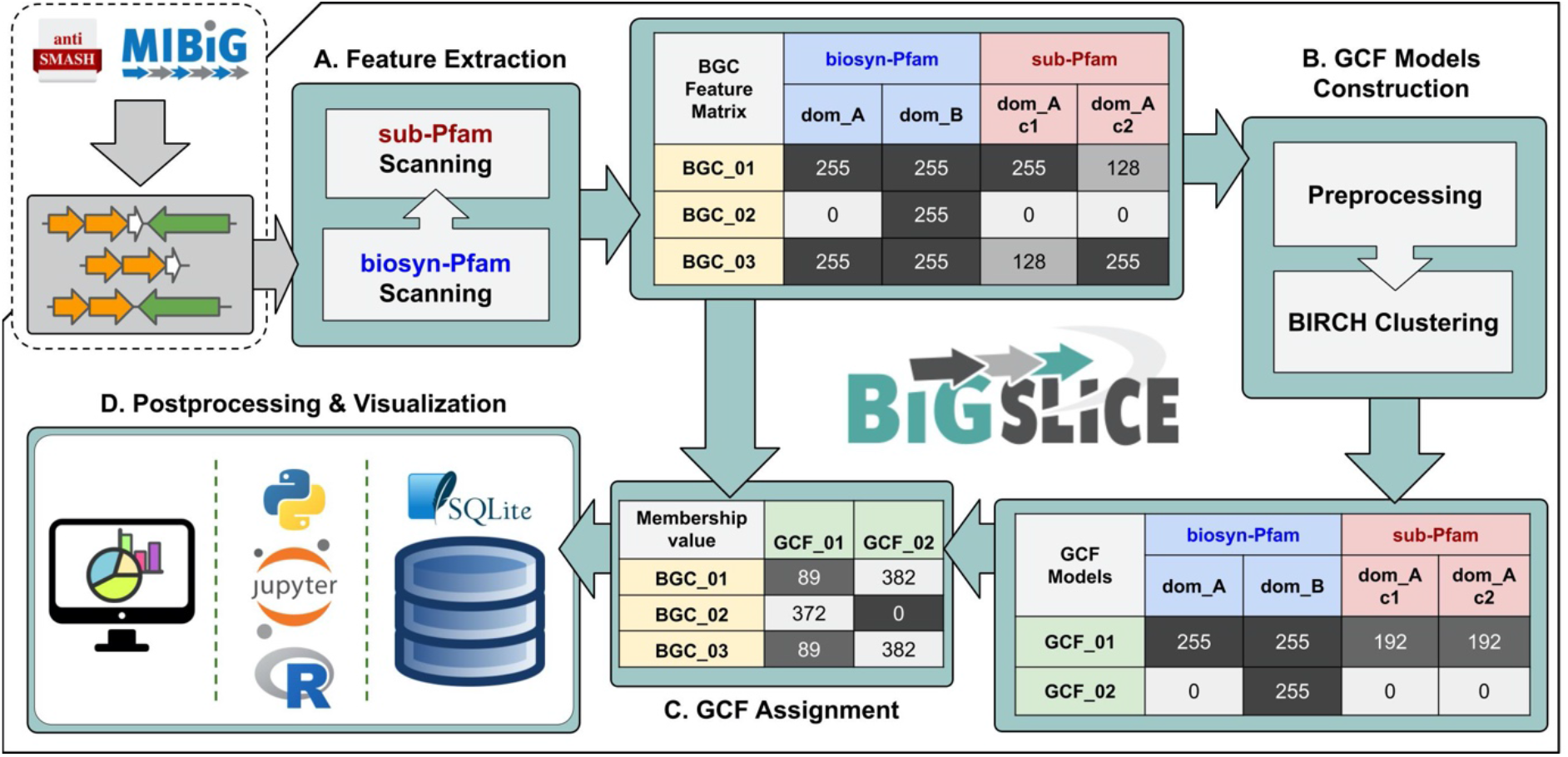
An overview of BiG-SLiCE’s GCF analysis workflow. Taking an input of region/cluster GenBank files from antiSMASH and MIBiG, **A.** BiG-SLiCE converts BGCs into numerical feature vectors, which are used to **B.** construct the GCF models (cluster centroids) and **C.** calculate BGC-to-GCF membership values. Processed data and results are all stored in a file-based SQL database (using SQLite3 [38]), which can then be used **D.** to perform further analysis (via external scripts) or to visualize the result in a user-interactive application.

### BGC feature extraction

In BiG-SCAPE, the (shared) occurrence and synteny (order) of Pfam [39] domains is measured for each pair of BGCs, along with the sequence similarity of homologous core genes, in order to construct a pairwise-distance network and define GCFs in this network using the Affinity Propagation algorithm [40]. While this hierarchical approach enables a very sensitive measurement of the relationships between BGCs and provides networks that can be interactively explored, it leads to a quadratic runtime complexity that does not allow application beyond a few tens of thousands of BGCs. To enable more efficient calculation of GCFs via partitional, near-linear time complexity clustering algorithms such as K-means [41] or BIRCH [42], we need to transform BGCs into numerical feature vectors (commonly known as quantization). We do this using a combination of two approaches: 1) biosynthetic domain (biosynthetic-pfam) absence/presence matrix construction and 2) signature domain (sub-Pfam) fingerprinting.

### Feature set 1: biosynthetic domain absence/presence matrix (biosynthetic-Pfam)

Domain hits (retrieved using hmmscan [43] with the gathering threshold) obtained for a reduced list of Pfam version 32 [39] pHMM models [Figure 2A] were used to construct a boolean (here represented by values of 0 or 255) feature matrix for every BGC. This list was constructed by filtering all Pfam domains for biosynthetically related protein families using the combination of ECDomainMiner [44] (which allows us to filter for domain related to enzymatic functions) and manual filtering based on each domain’s full description [Supplementary table 1]. This filtering was done to reduce the influence of non-biosynthetic domains, i.e. from genes that may be important for a BGC to function but are not directly responsible for generating structural variation of the produced metabolites (such as transporter enzymes and regulators). A library of 250 pHMM models from antiSMASH [19] was also included, as they harbor many curated biosynthetic domains not covered by the Pfam database alone. Altogether, this combination of 2,027 “biosynthetic-Pfam” models shows an increased selectivity compared to the full Pfam database when used to separate BGCs according to the chemical class of their predicted products [Figure 2C].

**Table 1.** Numbers of genomes and BGCs in all datasets included for the large scale diversity analysis. Numbers inside brackets indicate the total number of genomes assigned to each kingdom based on the subsequent taxonomy analysis. The “Others” category includes the kingdom of *Archaea*, *Viridiplantae* (from MIBiG dataset) and unassigned taxa. A complete list of all genome accessions and their BGC counts can be seen in [Supplementary table 3].

| Dataset Name | Study | Counts (Genomes, BGCs) |  |  |  |  |  |
| --- | --- | --- | --- | --- | --- | --- | --- |
|  |  | Bacterial |  | Fungal |  | Others |  |
| RefSeq complete bacteria | - | 19,169<br>(19,166) | 101,531 | 0<br>(0) | 0 | 0<br>(3) | 0 |
| RefSeq draft bacteria | - | 162,352<br>(162,297) | 959,061 | 0<br>(0) | 0 | 0<br>(55) | 346 |
| GenBank fungi | - | 0<br>(0) | 0 | 5,939<br>(5,905) | 123,816 | 0<br>(34) | 123 |
| GenBank archaea | - | 0<br>(1) | 2 | 0<br>(0) | 0 | 1,162<br>(1,161) | 2,109 |
| Parks 2017 (Uncultivated Bacteria and Archaea MAGs) | [54] | 7,280<br>(7,280) | 15,829 | 0<br>(0) | 0 | 623<br>(623) | 756 |
| Tully 2018 (TARA ocean MAGs) | [55] | 2,283<br>(2,326) | 4,829 | 0<br>(0) | 0 | 344<br>(301) | 518 |
| Almeida 2019 (Unified Human Gut MAGs) | [56] | 4,616<br>(4,616) | 4,766 | 0<br>(0) | 0 | 28<br>(28) | 25 |
| Stewart 2019 (Cow's rumen MAGs) | [57] | 4,815<br>(4,815) | 8,380 | 0<br>(0) | 0 | 126<br>(126) | 589 |
| Glendinning 2020 (Chicken's caecum MAGs) | [58] | 469<br>(469) | 481 | 0<br>(0) | 0 | 0<br>(0) | 0 |
| MIBiG v2.0 | [71] | 0<br>(0) | 1,594 | 0<br>(0) | 276 | 0<br>(0) | 40 |
| **Total** |  | 200,984<br>(200,970) | 1,096,473 | 5,939<br>(5,905) | 124,092 | 2,283<br>(2,331) | 4,506 |

**Figure 2.**
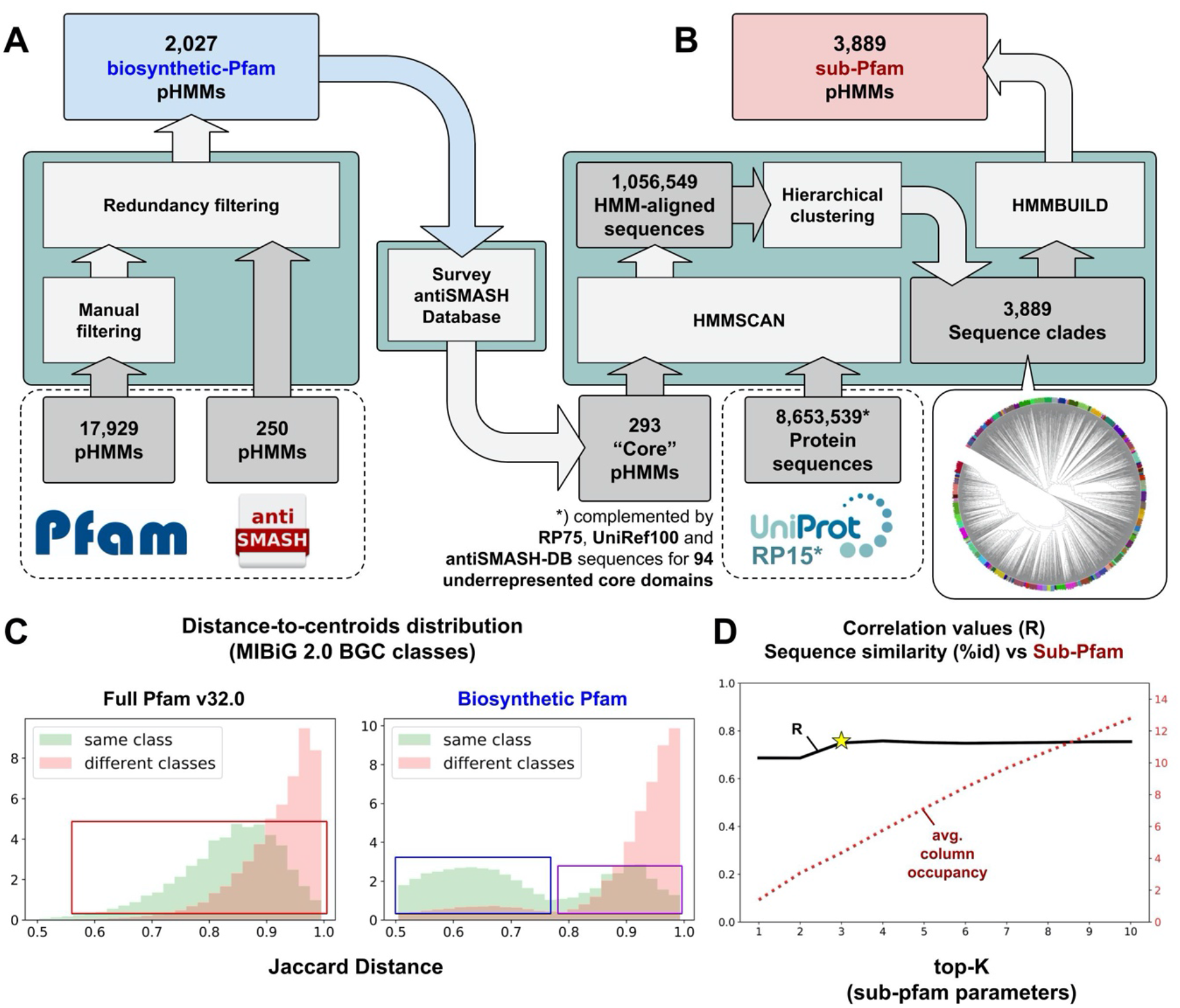
**A.** Construction of biosynthetic-Pfam features and **B.** Sub-level Pfam (sub-Pfam) features. **C.** Effect of Pfam model filtering on the discriminatory power of domain-presence Jaccard distance (JI index in BiG-SCAPE) measurements to separate MIBiG v2.0 generic classes (Polyketide, NRP, RiPP, Alkaloid, Terpene, Saccharide, Other). It is shown that the filtering strategy will produce more clearly separated within-class distances (blue box) than the full Pfam counterparts (red box). The second mode at the right side of the biosynthetic-Pfam same-class distribution (purple box) largely stems from hybrid BGCs, containing signature domains of two or more distinct classes (i.e. NRPS-PKS, PKS-Terpene-Saccharide, etc.). **D.** Pearson correlation values between protein sequence similarity (%-identity) and the corresponding sub-Pfam-based scoring in all AMP-binding domains (3,419 sequences, 879 BGCs) from the MIBiG v2.0 dataset across different top-K settings. Better correspondence (avg. R=0.75) is shown starting at top-K=3 (BiG-SLiCE’s default) onwards. The larger the top-K values, the more columns occupied (dashed red line) by the BGC’s composite sub-Pfam features as opposed to the biosynthetic-Pfam features, which can be thought of as a way to “tune” the core domain’s feature weight (akin to BiG-SCAPE’s anchor boost setting).

### Feature set 2: Signature domain fingerprinting (sub-Pfam)

While the biosynthetic-Pfam models work well to capture the pattern of BGC diversity across generic chemical classes, they are not sensitive enough to cover the more granular level of the inter-class diversity. BGCs of the same class typically share a limited set of “core” enzymes that determines the end product’s scaffold based on the combination of their specificity and/or copy number variation. For example, the compound’s scaffold produced by a Type-I Polyketide BGC is largely driven by the specificity of its (often multiple) Acyltransferase (AT) and Ketosynthase (KS) domains [45]. To cover this sequence-level protein diversity, we constructed alignments of 9,451,490 representative protein sequences in the RP15 database (Release 2020_01) [46] to our pre-selected 293 core biosynthetic domain pHMMs [Supplementary table 2]. We performed hierarchical clustering analysis to group similar aligned sequences into clades, then built sub-level protein family pHMMs from the sequences of each clade [Figure 2B]. This approach resulted in a distinct set of 3,889 sub-level Pfam (sub-Pfam) models (10-100 clades per core domain). For each aligned core domain in a BGC, an hmmscan search is performed using the specific sub-Pfam models, of which the hits are then ranked according to their bitscores. A set number of top hits (top-K) is then used to assign descending values of the corresponding feature in the matrix - for example, if a domain A has top-3 hits of A-c15, A-c3, and A-c2, its ranked feature values could be A-c15=255, A-c3=170 (255 x ⅔), and A-c2=85 (255 x ⅓). When a BGC has multiple hits on the same sub-Pfam column, the maximum value for that column will be taken. Using this ranked normalization scoring strategy for building the numerical feature representation of each core gene, we show that the sub-Pfams can together act as a proxy for sequence-level protein diversity [Figure 2D].

### GCF models construction

To efficiently group BGC features into GCFs, BiG-SLiCE uses a clustering method based on the python scikit-learn [47] implementation of the BIRCH [42] algorithm. When using gene cluster GBK files from antiSMASH v4.2 or higher (the version in which the attribute “on_contig_edge” was implemented to indicate which BGCs lie on the edge of a contig and may therefore be incomplete), users can opt to build the GCF features only from non-fragmented BGCs (using “--complete” parameter). Then, a distance sampling test will be performed to ascertain a default threshold value *T* for the clustering algorithm, unless a value is directly supplied by users via the “--threshold” parameter. The former is done by taking the average *Xth*-percentile (default *X*=1) of Euclidean pairwise distances between 100×1000 randomly sampled features from the input data. Afterwards, a flat-tree BIRCH (*branching_factor* >= *n_samples*) [48] clustering method is used to incrementally scan BGC features and build the GCF centroids. Then, a global cluster assignment is performed to match all input BGCs with the top-*N* (default *N*=3) scoring GCFs per BGC along with their membership scores. By considering multiple GCFs at once, users will be able to judge the confidence level of each BGC-to-GCF assignment. This is useful, for example, when determining the context of a fragmented BGC, where (low) membership scores might be distributed almost equally across different best-matching GCF models. Furthermore, by performing feature extraction on a set of newly sequenced (putative) BGCs, users can immediately match them with previously calculated GCF models (using the “--query” mode of BiG-SLiCE) and retrieve information on their characteristics and potential novelty.

### Comparison against manually curated GCFs

In order to judge the quality of results produced by its heuristic-based algorithm, we compared BiG-SLiCE clustering against 92 manually curated groups of MIBiG v1.3 BGCs provided in the original BiG-SCAPE paper. Several different threshold parameters *T* were tested (300 - 1,500) and corresponding results were compared to the reference groups. We calculated the V-score [49] of each run, which measures both the homogeneity (whether cluster members share the same target class) and completeness (whether members from a single target group are assigned into exactly one cluster) of a clustering result when matched to a (manually defined) target reference [Figure 3A], and plot it alongside the difference of GCF counts (*Δ*GCF) between the two. We found that BiG-SLiCE produces a generally agreeable result at the selected “optimal” threshold (V-score = 0.81 on *T* = 1,100), but is not able to capture the “perfect” clustering denoted by the reference groups [Figure 3B]. This stems from the fact that the (manual) categorization of the 92 compound groups does not always translate into the groups sharing a similar distance distribution in the BGC space, making it impossible to set a single clustering threshold that reproduces the membership assignment. BiG-SCAPE seems able to handle this issue better (V-score = 0.91 [Supplementary Figure 1]) due to its Affinity Propagation [40] based clustering algorithm that allows finding non-convex clusters, as opposed to the spherical partitioning approach of BIRCH, which is one of the main trade-offs for its hyper-scalability. BiG-SLiCE however accurately captures the underlying biosynthetic signal that connects the genomic space of BGCs and the chemical space of their products, as demonstrated by the bimodal distribution of distances between BGCs within vs. between the curated groups [Figure 3C] and the visualized feature heatmap of the most challenging groups [Figure 3D].

**Figure 3.**
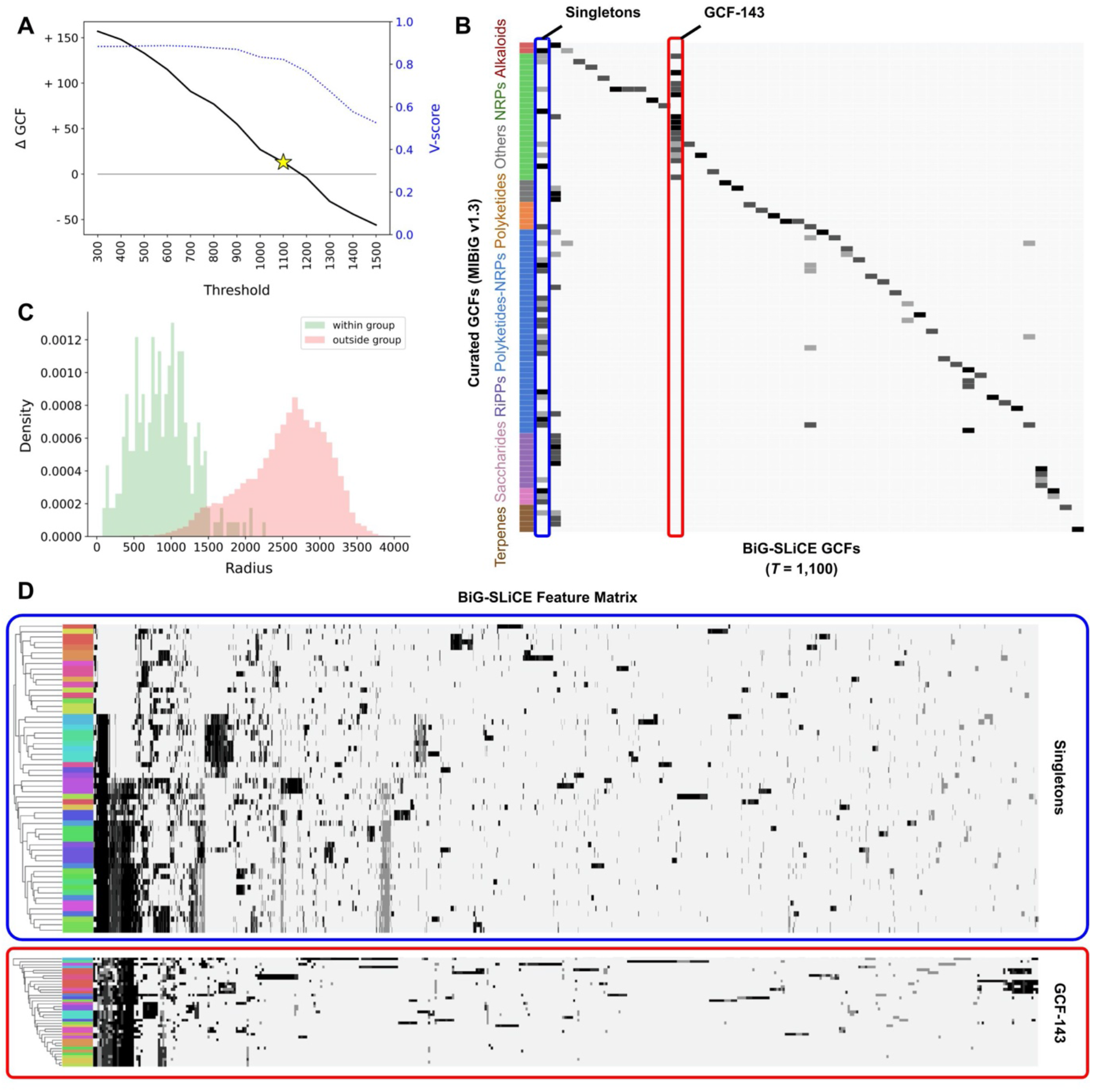
**A.** BiG-SLiCE analysis results for a range of threshold values, as measured by the difference of GCF counts (*Δ*GCF) and the level of clustering agreement (V-score of 1.0 for perfect clustering) compared to MIBiG curated groups. A single threshold result with the lowest *Δ*GCF while maintaining a V-score > 0.8, *T* = 1,100, was taken for further analysis in this figure. **B.** Confusion matrix of BiG-SLiCE clusters vs curated GCFs. To help in visualization, all singletons of the BiG-SLiCE result (58 GCFs) were collapsed into a single column (leftmost column, highlighted in blue box), showing together BGCs requiring a more lenient threshold (*T* > 1,100) to match the curated information. Conversely, another column, GCF-143 (red box), highlights the need for a stricter threshold (T < 1,100) to obtain a more fine-grained clustering for some parts of sequence space. **C.** BGC-to-centroid distance value (i.e. radius) distribution of within and between group pairs in the curated dataset. The centroid of each curated group was calculated by averaging the feature vectors of all BGCs assigned to it. **D.** Feature heatmap of the collapsed singleton group and GCF-143. Colored bars on the left indicate manually curated groups. In both cases, hierarchical clustering analysis (Euclidean-based, average-linkage) shows that the underlying pattern captured by BiG-SLiCE features tends to agree with the manually curated information, i.e. rows with the same color tend to be located near each other.

### SQL-based data storage enables extensive functionality

A typical BiG-SLiCE run produces a large amount of useful information on top of the GCF membership for each BGC. Taxonomic metadata, information on chemical compound classes and protein annotations are commonly included in the antiSMASH-generated BGC genbank files. To integrate that information and provide a truly comprehensive analysis output, a structured approach to data storage and processing is required. The architecture of BiG-SLiCE is centered around the use of a relational SQL database schema [Supplementary Figure 2] implemented as a file-based SQLite data store [38]. Processed input (including all metadata), supporting data and clustering results are systematically stored in the database tables. Using this setup, it is possible to build complex queries and perform all sorts of analyses even beyond the scope of GCF reconstruction. For example, one can use the preprocessed SQL database as a personal “data management” solution for custom BGC collections, enabling a fast search and query of specific protein sequences based on taxonomy and domain contents [Figure 4A]. Furthermore, this structured information about BGCs, their homology (GCF membership), taxonomy, biosynthetic classes, and protein domain hits can also be combined with a bioinformatics pipeline or analytical scripts written in Python or R (both of which have native support for SQLite) [Figure 4B] to perform even more complex analyses, for example to study the diversity of biosynthetic domains across samples and across taxonomy [Figure 4C]. As a matter of fact, all analyses performed in this study (see Results & Discussion) heavily benefitted from (and relied on) the data-wrangling convenience provided by BiG-SLiCE’s SQLite database.

**Figure 4.**
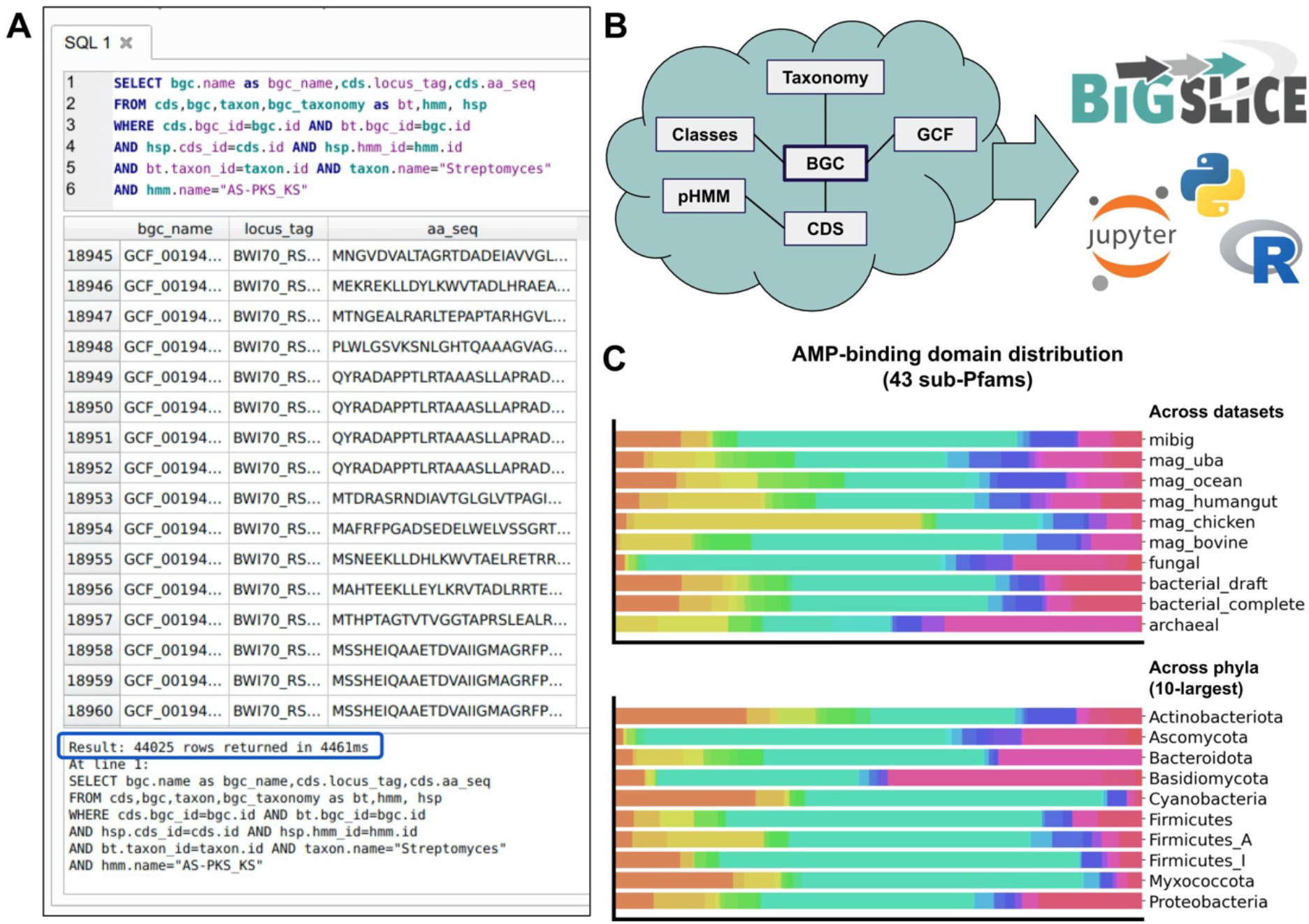
**A.** An example SQL query for all protein sequences harboring at least one Ketosynthase (AS-PKS_KS) domain from streptomycete BGCs. Here, the search performed against the total of ~29 million CDSes and >101 million domain hits in the database was completed in under five seconds, returning 44,025 CDS that satisfy the criteria. **B.** A cartoon illustration on how the interconnected SQL tables holding various BGC-related information can be leveraged by downstream analyses, e.g. using programs and notebooks written Python and R. **C.** An example downstream analysis using the data on sub-Pfam hits to chart the diversity of AMP-binding domains across datasets and across phyla. Here, each colored bar represents the distribution of a specific sub-Pfam clade across the sampled dataset / phylum. Each analysis including the SQL query took around 55 seconds to complete. A script to perform such analyses (which can also be used to investigate other biosynthetic domains) and generate the plots can be found in the “figure_4” folder of the Supplementary Data.

Finally, as previously demonstrated by the success of antiSMASH and BiG-SCAPE, one way in which regular end users can really benefit from a tool is when they are provided with an interactive and easy-to-use output visualization as a way to explore the data and analysis results. BiG-SLiCE offers this functionality by combining the portability of SQLite database with a mini web application written using Python’s Flask library [50]. This allowed us to implement a feature-rich visualization “software” that can be deployed and run with minimal amount of installation effort on a user’s personal computer (https://github.com/medema-group/bigslice#user-interactive-output). While this feature is currently at a prototype stage, offering simple functionalities such as browsing and viewing the processed BGCs and GCFs, we plan to continue to improve and implement more advanced features along the way, such as searching and filtering for specific BGCs / GCFs of interest, generating phylogenomic alignments of BGCs [22,51], or even incorporating additional useful information such as the presence/absence of antibiotic-resistant genes [52] and regulatory domains [53] within the BGCs.

## Results and Discussion

In order to show how BiG-SLiCE could be applied to large datasets that capture the full diversity of BGCs from cultured and uncultured microbes, we decided to collect a merged dataset of publicly available microbial genomes and metagenome-assembled genomes (MAGs). We then predicted their BGCs using antiSMASH v5.1.1, filtering out contigs < 5,000bp (“--minlength 5000”) and used the respective taxonomy options wherever applicable (“--taxon bacteria” for bacterial and archaeal genomes, and “--taxon fungi” for fungal ones).

### Collecting a near-comprehensive dataset of publicly available BGCs

We downloaded 19,169 complete and chromosome-level bacterial NCBI RefSeq genomes up to 27 March 2020, 12:15PM CET. To capture the extensive strain-level diversity within the bacterial kingdom, 162,352 draft RefSeq genomes were also downloaded and processed, resulting in a total number of 1,060,594 BGCs when combined. For fungi and archaea, we downloaded 5,939 and 1,162 genomes from NCBI Genbank with “Refseq-like” filters turned on, resulting in 123,939 fungal and 2,578 archaeal BGCs, respectively (all NCBI query scripts used for this data collection step are available in [Supplementary text 1]). Furthermore, we collected and processed 20,584 MAGs from previously published studies [54–58], resulting in a total of 36,173 BGCs. This list was arbitrarily selected from available studies describing the construction of large-scale MAG assemblies from different environments at the time of data collection. Although this list was in no way comprehensive (for example, there are many other notable recent publications [59–70] not covered by this initial effort, not to mention the huge number of shotgun metagenomic studies publishing only contig-level assembly of unassigned bins), the ~20K MAGs presented here may already give us a glimpse on the untapped biosynthetic diversity of uncultured microbes. Finally, we incorporated all 1,910 entries from MIBiG v2.0 [71] as a reference set of known and experimentally verified BGCs. In total, a final count of 1,225,071 BGCs were predicted from 209,206 genomes and MAGs, as shown in Table 1.

### Improving the taxonomy assignment of genomes

Before performing any taxonomy-related diversity analysis, we ensured that all included genomes were correctly assigned to their respective taxa. Several studies pointed out that there might be a potentially widespread misclassification of bacterial genomes within the NCBI database [72–74]. To avoid this issue, we chose to use the taxonomy derived from the GTDB (Genome Taxonomy Database), which were posited to be more phylogenomically accurate than that of NCBI [75]. We queried all bacterial and archaeal NCBI genome accessions through the GTDB API (version 04-RS89, https://gtdb.ecogenomic.org/api/) to fetch their taxonomy information, resulting in 123,245 taxonomy-assigned genomes. For the remaining genomes, i.e. those from metagenomic studies and more recent NCBI genomes not yet covered by the API, we used the GTDB toolkit [75], a bioinformatics pipeline that integrates several tools [43,76–80], to infer their taxonomy based on their genomic marker composition. This further assigned taxonomy information to another 79,964 genomes. Original NCBI taxonomy information was retained for all fungal genomes and MIBiG BGCs (a list of all GTDB and NCBI-assigned taxonomy per genome is available in [Supplementary table 3]).

### Large-scale Homology Analysis of 1.2 Million BGCs

We then performed BiG-SLiCE clustering analyses over the merged datasets using a 36-core, 252GB RAM shared computing server facility. Taking advantage of the antiSMASH5-enabled annotation of fragmented BGCs (clusters residing on contig edges), the “--complete-only” parameter was used for the clustering phase, using 802,287 (65%) non-fragmented BGCs from the input data to build the GCF models. This ensures that the variation in the models is derived from actual BGC diversity and not due to technical gene losses (from contig splits). Later on, the full input datasets were queried back against the GCF models, in order to map the fragmented BGCs onto their corresponding GCFs based on the calculated membership values *d*. For this analysis, we arbitrarily categorize GCF-to-BGC relationships into “core” (*d* <= *T*), “putative” (*T* < *d* <= 2*T*), or “orphan” (*d* > 2*T*) on a best-hit basis (parameter --n_ranks=1). Five different threshold values (*T* = {300, 600, 900, 1,200, 1,500}) were tested, producing a decreasing number of GCF models (more BGCs per GCF) as *T* gets bigger (more lenient) [Supplementary table 4]. The first run (*T* = 300) which carries the full workflow load (from features extraction to membership assignment) was finished in ~240 hours (10 days), or >150x faster than the estimated runtime of BiG-SCAPE [Supplementary Figure 3]. A large chunk of this runtime is spent at the feature extraction step, which includes the I/O heavy hmmscan and non-parallelizable SQL inserts [Figure 5A]. Subsequent runs (*T* = 600 - 1,500) reused the precalculated features, taking only an average of ~4 hours runtime for each run [Figure 5B].

**Figure 5.**
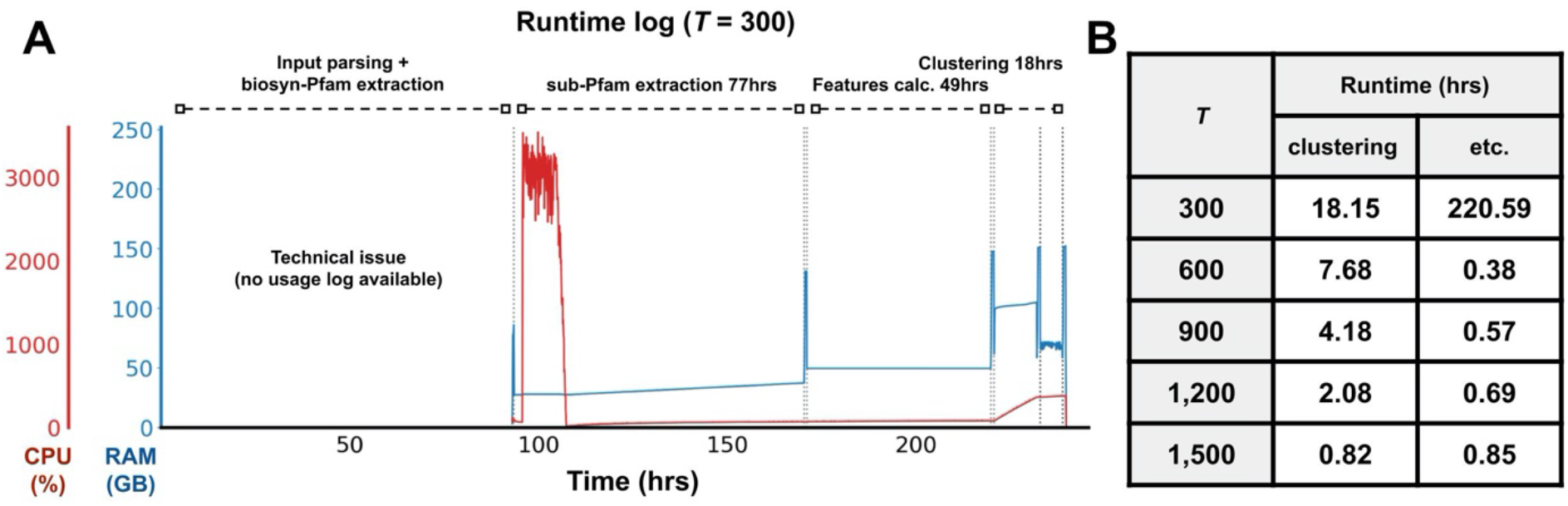
**A.** Runtime breakdown of the full run (*T* = 300) on a 36-core CPU, 262GB RAM server. Due to some technical issues, no usage log is available for steps prior to the sub-Pfam extraction. CPU usage log shows that most of the time, BiG-SLiCE only uses one CPU core, giving a room for further improvement e.g. via SQL parallelization. Spikes on the RAM usage (peak = ~150GB) came from the periodic “dumping” of the in-memory database (used in order to speed up runtime) into an SQLite db file. **B.** Runtime comparison between multiple runs, with *T* = 300 bearing the full load of performing input processing and features extraction. Here, runtimes are separately shown for both the clustering (GCF models construction + membership assignment) and other steps (input parsing, hmmscanning and features extraction).

### Charting a global map of BGC diversity

Each GCF in the global clustering analysis result represents a functional niche captured from a group of BGCs sharing a similar biosynthetic make-up. To enable the visualization of this biosynthetic diversity, we partitioned the 121,299 centroid features of the GCFs produced by the *T* = 300 run into 500 GCF “bins” using K-Means (via sci-kit’s library, with *K* = 500 and a random, but reproducible initialization step; see the reproduction script included in Supplementary Data for details). Another round of membership assignment was performed to match the full set of 1.2M BGC features into the resulting 500 GCF bin centroids. Those centroids were also subjected to an average-linkage agglomerative clustering analysis (sci-kit implementation, euclidean distance). The produced hierarchical tree object was then converted to a newick file (using a custom script provided in the Supplementary Data) and plotted via the iTOL web server (https://itol.embl.de/) [81]. By annotating this tree with various types of quantitative information [Supplementary table 5], the resulting phylogram pictures a generic, “bird-eye view” on the entire set of 1.2 million BGCs [Figure 6].

**Figure 6.**
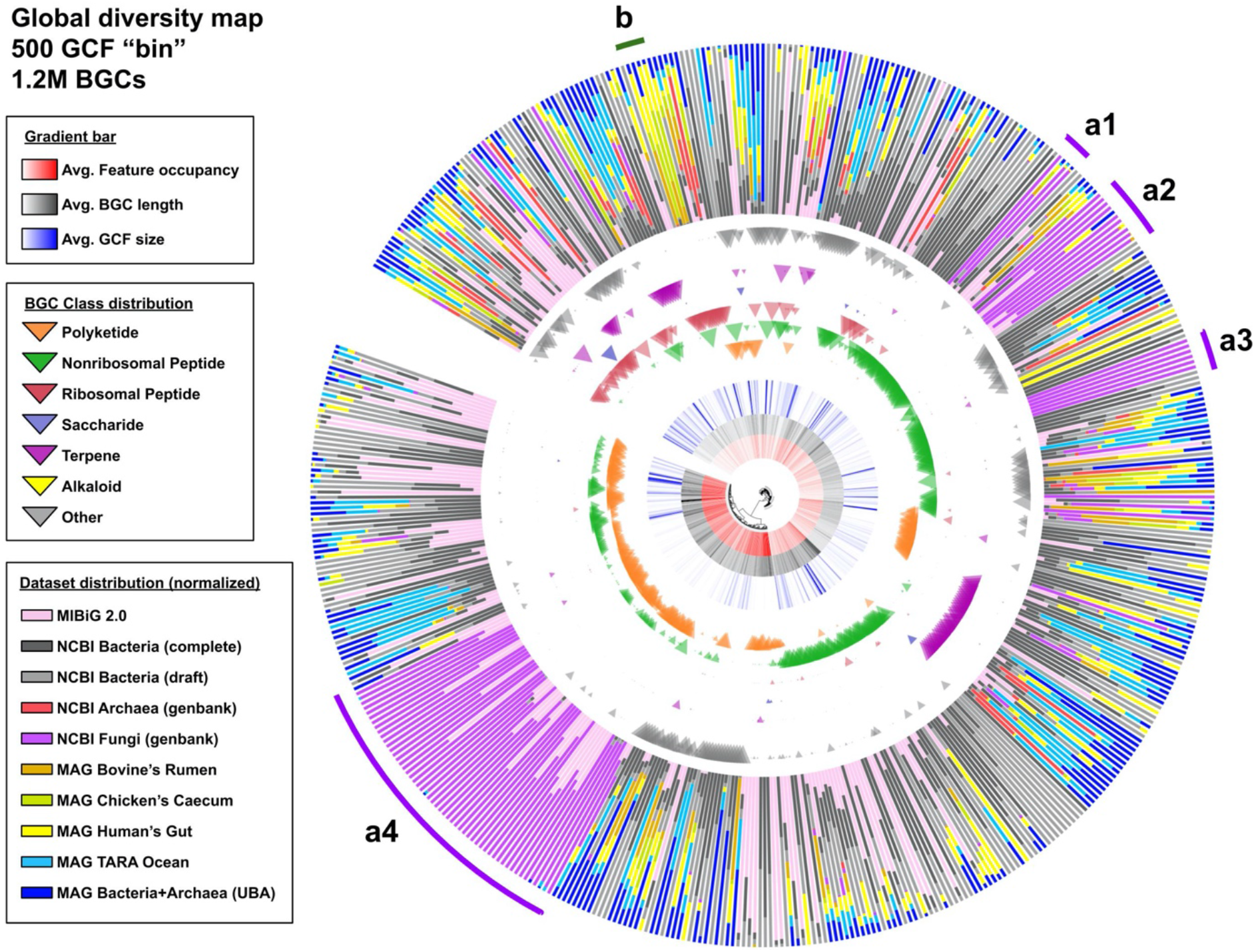
A phylogram created via the hierarchical clustering analysis of 500 GCF bins. The phylogram was rooted on a null (all zeros) dummy feature matrix. For each node, the raw dataset distribution values [Supplementary table 5] were double-normalized, first against the number of BGCs each dataset has in total, giving the fraction values, then against all fraction values of other datasets in the bin. Furthermore, some notably interesting clades were manually highlighted (a1-a4, b) for follow up discussion (see main text).

An important thing to note is that due to the non-deterministic nature of K-means, the number of BGCs that goes into each bin depends a lot on the randomly placed initial centroids (for example, there are 21 bins made up of a single BGC [Supplementary table 5], which can happen when the randomly placed initial centroid hits an outlier/singleton in the dataset). This is analogous to taking a two-dimensional satellite picture of the earth from a specific coordinate, looking down at a specific angle. There are an infinite number of ways to take a picture, giving a different perspective and snapshot of an object each time, but the inherent three-dimensional structure of the object will always remain constant. While the map shown in [Figure 6] can give us insights into the major “landmarks” formed by the larger groups of BGCs, it will not show all the nooks and crannies to be explored from the entire dataset (which could be explored using more fine-grained tools such as BiG-SCAPE).

The very first thing that we can notice from the phylogram is how fungal BGCs (purple bars, “a1” to “a4”) have quite distinct features that discriminate them from the rest of the (mostly bacterial) datasets. Clades “a1” to “a3” contain mostly NRP (99.93%) BGCs: 20,398 from “a1”, 18,770 from “a2” and 8,606 from “a3”. Clade “a1” shares its 9,402 fungal BGCs with 10,972 bacterial (67.56% came from *Pseudomonas*) and 13 archaeal ones. This clade includes two simple NRP-encoding fungal BGCs from MIBiG dataset, encoding the biosynthesis of the proteasome inhibitor fellutamide B [82] (BGC0001399) and aspergillic acid [83] (BGC0001516) from *Aspergillus* (and on the bacterial side: four MIBiG BGCs including another simple proteasome inhibitor livipeptin [84,85] encoded by BGC0001168 from *Streptomyces lividans*). Clade “a2” contains a major part (50 out of 61) of known non-hybrid fungal NRP BGCs in MIBiG, and shares the clade with 85 bacterial NRPs. Last but not least, clade “a3” almost exclusively (except for 1 beta-lactam BGC from *Mycobacterium gordonae* and 10 BGCs from unknown taxa) consists of uncharacterized fungal NRPs. A closer look at this clade leads to an interesting observation in terms of shared features / domains. We found that no domain (even at biosynthetic-Pfam level) is shared by more than 70% of the BGCs, except from a few sub-Pfams: AS-NAD_binding_4-c7 (91.92%), AS-AMP-binding-c6 (98.84%) and Epimerase-c26 (99.03%). These domains are often contained in one protein-coding gene, sometimes with an extra ACP (AS-PP-binding) domain (found in 75.34% of the BGCs). This clade therefore seems to contain mostly proteins related to α-aminoadipate reductases, which have been previously inferred to have an evolutionary origin prior to, or early in, the evolution of fungi [86]. Detailed results and reproducible scripts for analyses from this and subsequent paragraphs can be found in the “figure_6+sup_table_5” folder of the Supplementary Data.

At the opposite side of the phylogram, 42,716 out of 43,840 (97.43%) BGCs from clade “a4” are of the Type-I Polyketide (T1-PKS) subclass, and as many as 7,811 of them are “true” PK/NRP hybrids (determined by the presence of Acyltransferase, Ketosynthase, AMP-binding and Condensation domains together in the BGC). This clade shows an enrichment of AS-PKS_AT-c7 (95.1%) and ketoacyl-synt-c8 (95.94%) sub-Pfam domains possibly linked to the iterative mechanism almost exclusively attributed to fungal PKSes [87]. Interestingly, 2,255 BGCs from this clade have bacterial origins (966 mycobacterium, 438 streptomyces, 851 others), which might possibly be connected to a group of non-canonical, iterative T1-PKSes from bacteria [88–90]. However, no bacterial BGC from MIBiG, including those of known iterative type [91,92], falls into this clade.

We can also see a narrow but distinct clade “b” highly represented by RiPP BGCs from the “gut” metagenome datasets (bovine’s rumen, chicken’s caecum, human gut). Aside from the 2,546 (17.88% of the three datasets total) MAG-derived BGCs, this clade also contains 4,254 BGCs from the NCBI bacterial RefSeq genomes (0.40% of the dataset’s total) and is populated by BGCs from various kinds of firmicutes (99.32% of the clade’s total). Looking closer at the BGC classes gives away an important clue: 99.68% of the BGCs belong to the sactipeptide RiPP subclass as annotated by antiSMASH, and seems to encode a group of RiPPs known as SCIFF (Six-Cysteine in Forty-Five) peptides [93] (recently proposed to be reclassified as ranthipeptides [94]), as 100% of those RiPPs have the signature TIGR03973 precursor domain (along with >99% occurrence of Radical_SAM and the iron-sulfur binding Fer4_12 domains). It is largely unknown why this particular class of BGCs are highly represented in the gut microbiomes, except for the fact that they can only be found in typical resident microbes of those environments (80.52% of BGCs came from *Clostridia*). Recently, a series of analyses performed by Chen et al. in solventogenic *Clostridia [95]* suggested that these RiPPs might play a role in the quorum sensing system and in controlling cell metabolism of such organisms.

Next, by looking at how the pink (innermost) bar is spread all across the phylogram, we can infer that despite holding no more than 2,000 entries presently, the BGCs in the MIBiG database are actually diverse enough to cover much of the general diversity of BGCs. However, we also need to be aware of the fact that most of the detection rules in antiSMASH were almost directly derived from the knowledge of experimentally characterized BGCs that are also present in MIBiG. This means that the 1.2 million BGCs we captured from those 209 thousand genomes are all evolutionarily related, although distantly, to at least one MIBiG BGC. To go beyond these canonical pathways, several unsupervised but “lower-confidence” alternative algorithms [24,96] have been developed that can potentially complement antiSMASH to cover more exotic areas of biosynthetic space.

Finally, this visualization suggests that several aspects can still be improved upon this first version of the BiG-SLiCE clustering algorithm. The three innermost gradient bars of the phylogram show the variation in the length of BGCs, extracted features, and the size of GCFs. By looking at them, it is quite apparent that there is a distinct separation between two major groups of GCF bins: a high feature counts group (more intense red bars) consisting mostly of domain-rich Polyketide (and some nonribosomal peptide / NRP) BGCs, and a low feature counts group (less intense red bars) consisting a large majority of NRP BGCs along with most Terpene and RiPP BGCs [Supplementary figure 4A]. This causes a large dichotomy in GCF sizes [Supplementary figure 4B] due to the limitation of the single-threshold clustering method of BIRCH as described before. While, generally, the number of extracted features depends a lot on the length of a BGC (longer BGCs may contain more genes and domains), this is not always the case. For example, there may be a great degree of copy number variation between biosynthetic domains (e.g. in some NRP BGCs) that is not captured by BiG-SLiCE [Supplementary figure 4C], as it only looks at absence/presence patterns of (sub-)Pfam features. Additionally, the pHMM models of BiG-SLiCE may fail to capture the diversity of certain tailoring domains. Conversely, there are also cases where the structure of the end products depends largely on the residue-level variability of particular proteins, such as for the large majority of RiPP BGCs, in which biochemical variation is largely governed by the sequences of precursor peptides [Supplementary figure 4D]. Thus, one way to optimize BiG-SLiCE clustering in the future is to try and balance the average feature counts across BGC (sub)classes, i.e. by surveying and including the missed neighboring domains, by putting more emphasis on core domain specificity (more columns for subpfam models) of a manually selected set of enzymes, and/or by taking into account copy number variation of domains (e.g. counting the actual number of biosynthetic-pfam hits rather than using a boolean absence/presence value). Alternatively, large BiG-SLiCE GCFs can be analyzed in more detail using BiG-SCAPE or using protein sequence similarity networks [97] (which can, for example, be very powerful for analyzing RiPP precursor peptide variation [98–100]).

### Measuring the “hidden iceberg” of microbial secondary metabolism

Only limited numbers of studies have considered global measurements of biosynthetic potential across taxa, or comparisons between cultivated and uncultivated bacteria [23,24,101,102]. To demonstrate how BiG-SLiCE could be used in such studies to quantify unexplored biosynthetic potentials, we took the 29,955 GCFs calculated from *T* = 900 and measured the distance of every GCF model against their closest MIBiG BGC features [Supplementary table 6], then plotted a histogram from the data [Figure 7A].

**Figure 7.**
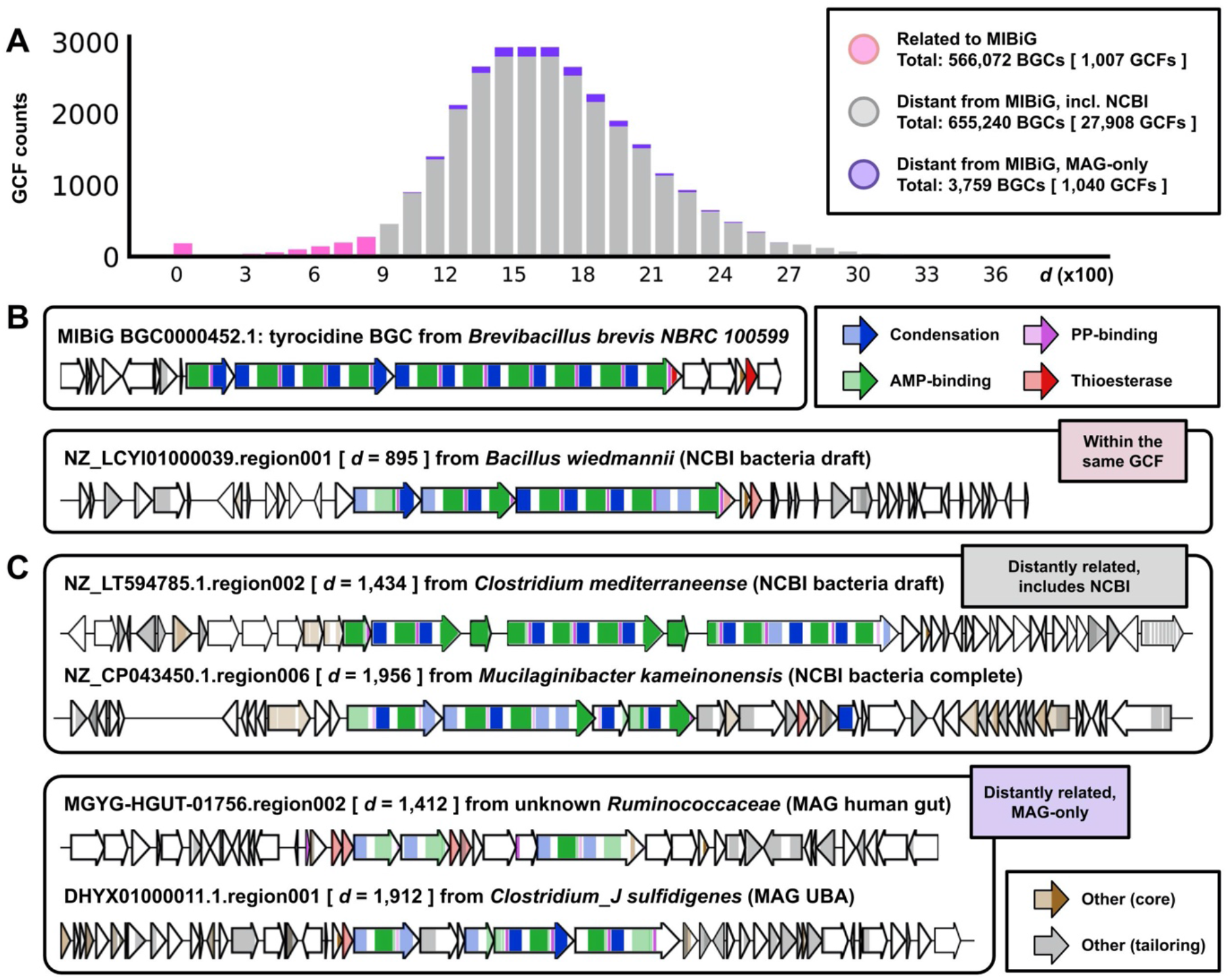
**A.** Histogram of Euclidean distances (x-axis) of GCF models to their closest BGC from the MIBiG 2.0 dataset. Here, all GCFs having *d* <= 900 were denoted as “Related to MIBiG” and “Distant from MIBiG” if otherwise, particularly highlighting those coming only from the MAG datasets. **B.** Selected anecdotal example of a MIBiG BGC and one of the farthest (*d* = 895) BGCs from the same GCF, which does not encode a biosynthetically equivalent pathway. Colored sections of the arrows represent biosynthetic domains captured by BiG-SLiCE, where darker colors represent putative core domain homologues (as measured by the sub-pfam signature) shared between the MIBiG BGC and its distant relatives. **C.** Example BGCs from GCFs having a distant best-hit to the tyrocidine BGC as shown by their generally high *d* values (1,412 - 1,956) to the MIBiG BGC in question.

Indeed, it is immediately clear from [Figure 7A] that the great majority (96.63%) of GCFs remain uncharacterized (distantly related to any MIBiG BGC), representing a huge iceberg of unknown secondary metabolism hidden under the surface represented by the MIBiG database. Of these 28,948 GCFs, 1,040 can only be found in MAG datasets, representing unique BGCs from uncultured and unculturable microbes. However, care should be taken not to accept the numbers at face value, as there are still a lot of factors yet to be considered. On the one hand, while we previously showed that the 1,910 BGCs in MIBiG have good diversity coverage across biosynthetic classes, the database is not entirely comprehensive in capturing all experimentally characterized BGCs to date. On the other hand, the arbitrary threshold used to define the relationship (*T* = 900) might be too lenient in some cases, as shown by an NRP BGC seemingly unrelated to the tyrocidine BGC being put together in the same GCF [Figure 7B]. This also means that many BGCs with very low feature count would be lumped together in a large GCF with some MIBiG ones, contributing to an overestimated number (566,072 BGCs, or 46.2% of total input) of BGCs “related to MIBiG BGCs”. Combined with the fact that the analysis only includes what antiSMASH covers, we argue that the actual number of BGCs encoding distinct secondary metabolic pathways unrelated to known ones is likely to be even bigger.

### Exploring biosynthetic potential across taxonomy

One of the potential use cases of BiG-SLICE is the systematic exploration of biosynthetic potential across taxonomy, which may provide detailed insight to direct discovery efforts. Having the species information of 209,206 genomes at hand, we sought to showcase how such an application could work by calculating the total number of GCFs within species having four or more strain-level genomes from our datasets (a total of 3,181 species from 1,043 genera) [Supplementary table 7]. To get a rough idea on the alpha diversity of GCFs within each species, we used the result of two threshold parameters, *T* = 300 and *T* = 900, and counted the numbers of GCFs per species across the two runs [Figure 8A]. In this scenario, three *Firmicute*s (*Bacillus velezensis*, *Bacillus thuringiensis*, *Streptococcus pneumoniae*) and five *Proteobacteria* (*Escherichia flexneri*, *Klebsiella pneumoniae*, *Acinetobacter baumannii*, *Escherichia coli, Burkholderia ubonensis*) dropped out of the top-30 list of richest species when going from the stringent threshold to the more lenient one. This suggests that the perceived GCF richness in those species was largely confounded by the effect of (multiple) gene insertions/deletions near BGCs (in flanking regions included by antiSMASH) rather than the actual recruitment of new BGCs (i.e. via lateral gene transfer [103–105]).

**Figure 8.**
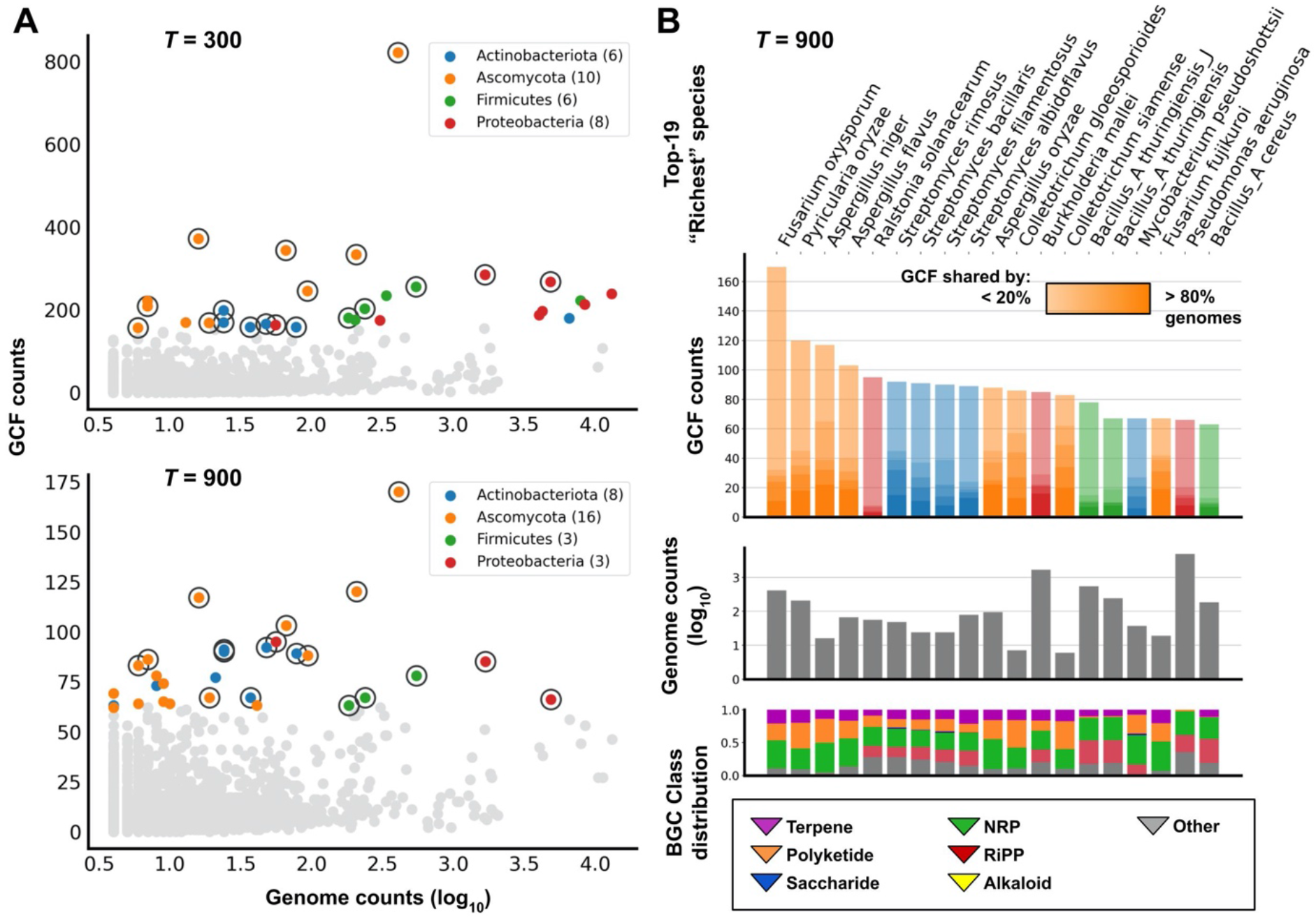
**A.** Distribution of GCF counts across species having four or more genomes in the dataset. Two plots showing results at the most stringent (*T* = 300) and a fairly lenient (*T* = 900) threshold, each highlighting 30 species with the highest GCF counts (colored dots). 19 species present in the top-30 of both thresholds are marked with black circle. **B.** Detailed view of the top-19 species, taking GCFs from the *T* = 900 result. Gradients from the colored bars (GCF counts) represent the extent to which a GCF is shared between all genomes in a species (in 20%-wide steps) [Supplementary table 7]. Additionally, the total distribution of BGC classes per species is also measured [Supplementary table 8].

Four *Streptomyces* species made it into the selected list of 19 species that consistently ranked top-30 in both runs [Figure 8B] despite having relatively few genomes (24 to 78) in the dataset, confirming their status as prolific producers of natural products: 75-80% of approved antibiotics are sourced from this genus alone [1,106]. More detailed analysis of the set of species that have precisely four genomes in the dataset (723 species from 486 genera) [Supplementary table 7] showed that 26 species (104 genomes) from this “run-of-the-mill” drug discovery genus harbor an average number of 36.69 unique GCFs (at *T* = 900) per species, putting it first among other bacteria, followed by *Saccharopolyspora* (36 GCFs from 1 species), *Nocardia* (avg. 30 GCFs from 2 species), and *Amycolatopsis* (avg. 29 GCFs from 3 species).

The rest of the bacterial species (1 actinobacterium, 3 firmicutes and 3 proteobacteria) that made it into the top-19 are mainly composed of pathogens that have had many of their genomes sequenced (183 to 4,838 genomes) within the NCBI database, which contributes greatly to their elevated GCF richness measure. However, two species from the list showed numbers that deviate from this observation. *Mycobacterium pseudoshottsii*, a slow-growing fish pathogen originally isolated from striped bass (*Morone saxatilis*) during mycobacterial outbreak in *Chesapeake Bay* [107] harbors a total of 67 unique GCFs within its 37 genomes. This makes the species distinct compared to the rest in the genus: *Mycobacterium avium* which harbors 58 GCFs from 197 genomes followed by *Mycobacterium tuberculosis* with 56 GCFs from 6,606 genomes. However, a closer look shows that the majority (35 out of 37) of the GTDB-Tk assigned genomes from this species actually belong to the closely related *Mycobacterium marinum* and *Mycobacterium ulcerans* in NCBI, which might explain the group’s observed higher total GCF diversity. These accessions are now included and are assigned correctly in the newer version of GTDB R05-RS95 (and the accompanying GTDB-Tk version 1.3.0).

*Ralstonia solanacearum* (also known as *Pseudomonas solanacearum*), the final pathogenic species from the bacterial list actually made it into the top-5 (first place among bacteria) with 95 GCFs derived from its 56 genomes. A striking observation from this species data is how little overlap occurred between the BGCs from different strains: 87 out of the 95 (91.5%) GCFs are shared only between less than 20% of strain genomes, meaning that every 11 strains may harbor ~17 unique BGCs that cannot be found in any other strain of the species. Not much can be said about the potential natural products that can be mined from this diversity (two hybrid NRP/Polyketide compounds, an antimycoplasma micacodin [108] and a fungi-colonizing agent ralsolamycin [109] from a tomato-associated strain GMI1000, were deposited in MIBiG under accessions BGC0001014 and BGC0001363/1754), but several comparative genomic analyses [110,111] have linked this highly divergent metabolic capacities with their unusual ability to attack a vast range of plant species [112].

Finally, fungal secondary metabolism presents an enigma in the space of natural product and drug discovery: although some of the most important drugs came from fungi, such as cyclosporine, penicillins and lovastatin, they arguably remain underexplored when compared to the bacteria. Indeed, there are only 88 entries from *Aspergillus* as opposed to 636 from *Streptomyces* in MIBiG 2.0. Similarly, there are around 2,000 streptomycete genomes in NCBI GenBank compared to ~400 from *Aspergillus*. This phenomenon might be attributed to the general difficulty of working with filamentous fungi, due to, e.g., their relatively complex genomes. Nevertheless, many fungal species managed to place themselves onto the list of species with the richest GCF repertoires. As many as 32 ascomycota from 17 different genera were part of the top-100 ranked species in the *T* = 900 list, and despite its lower genome count (410) compared to, e.g., the bacterial pathogen *Pseudomonas aeruginosa* (4,858), *Fusarium oxysporum* managed to top the chart with 821 unique GCFs. Similarly, three *Aspergillus* species have a genome-to-GCF ratio similar to, or in some cases higher than the *Streptomyces* species on the list. As fungi and bacteria seem to frequently compete with each other in the wild [113], it may be logical to increase the search for new antibacterial compounds from this nemesis of bacteria, complementary to bacterial genome mining.

### Conclusions and Future Perspectives

Here, we demonstrated that with BiG-SLiCE, we finally have the means to generate and exploit a truly global map of secondary metabolic diversity, which can provide insights for both fundamental (studying the diversity and evolution of microbial secondary metabolism) and practical (drug and novel compound discovery) purposes. To draw more solid biological conclusions from this kind of analysis, the issue of uneven feature coverage needs to be addressed (leading to some BGCs being more granularly clustered than others at any given threshold) and a more robust approach needs to be designed for choosing a threshold for clustering.

One important topic that has not been discussed extensively is how we can deal with fragmented BGCs. This is especially important when considering incorporation of more MAGs and shotgun metagenomic data in future analyses. Although the fuzzy membership approach provides a way for an objective (manual) inspection of BGC placement, an automatic but statistically-informed placement strategy still needs to be developed (as opposed to taking only the best hit coupled with some arbitrary thresholds as done here). Additionally, implementing a vector-based counterpart of BiG-SCAPE’s “glocal” comparison, which matches only the aligned fraction of a complete BGC against a fragmented one (e.g. by only calculating the euclidean distance of shared columns) might help to dampen the effect of the variable feature size each GCF had.

While this first version of the software constitutes a big leap in scalability of BGC analyses, a long road is still ahead. We invite the community to help improve BiG-SLiCE by sending feedback and using it to investigate the many specific questions that they have which were impossible or highly impractical to answer before. Finally, while a similar massive-scale BGC analysis can be performed *ad hoc* given sufficient computational resource and expertise, we plan to convert the precalculated global analysis result into a publicly accessible “reference” GCF database, allowing the scientific community to benefit from the result in new ways. For example, by curating this reference database with structural and functional annotations derived from (known) BGCs, it can facilitate the functional characterization and dereplication of newly sequenced BGCs.

## Availability of Supporting Data and Materials

Project name: BiG-SLiCE

Project home page: https://github.com/medema-group/bigslice

Operating system(s): Linux / UNIX-based OS, output web app can be viewed on any modern Internet browsers

Programming language: Python

Other requirements: Python 3.6 or higher License: GNU Affero General Public License v3.0

Input BGCs, analysis results and python scripts used to generate all figures and tables in this study is available at https://bioinformatics.nl/~kauts001/ltr/bigslice/paper_data/. An archived v1.0.0 release of the BiG-SLiCE software including the pHMM models used for this study can be downloaded from Zenodo [114].

## Supplementary texts

All supplementary texts are available via BioRxiv.

### Supplementary figures

**Supplementary Figure 1.**
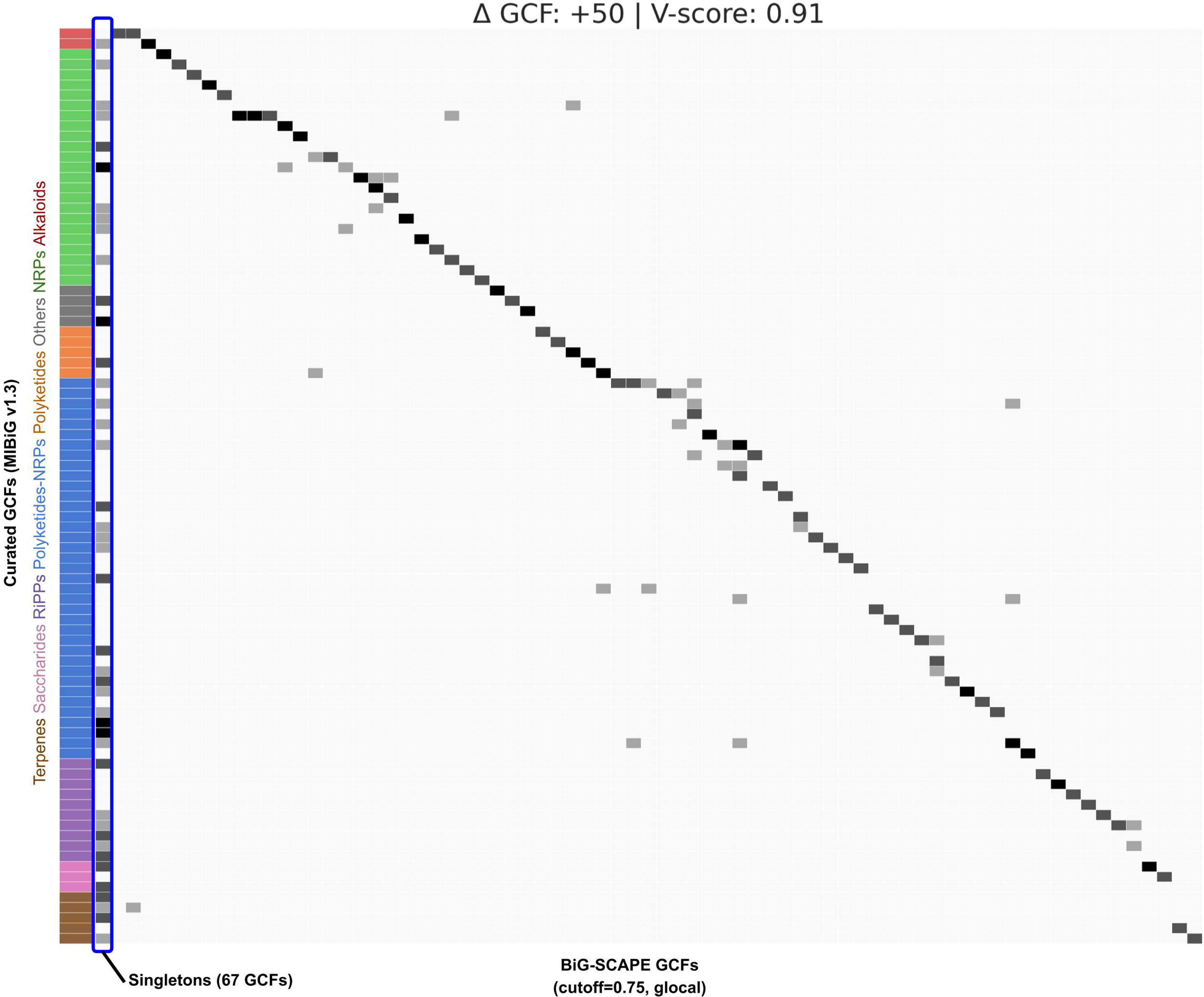
Confusion heatmap of BiG-SCAPE result compared to the curated set of MIBiG BGCs. The result was generated using BiG-SCAPE version 1.0.1, using a cutoff threshold of 0.75 and hybrid mode turned off, as specified in the original paper. A “vertical band” is highlighted in blue, comprising BGCs unintentionally assigned as singletons due to the strictness of the cutoff parameter being used.

**Supplementary Figure 2.**
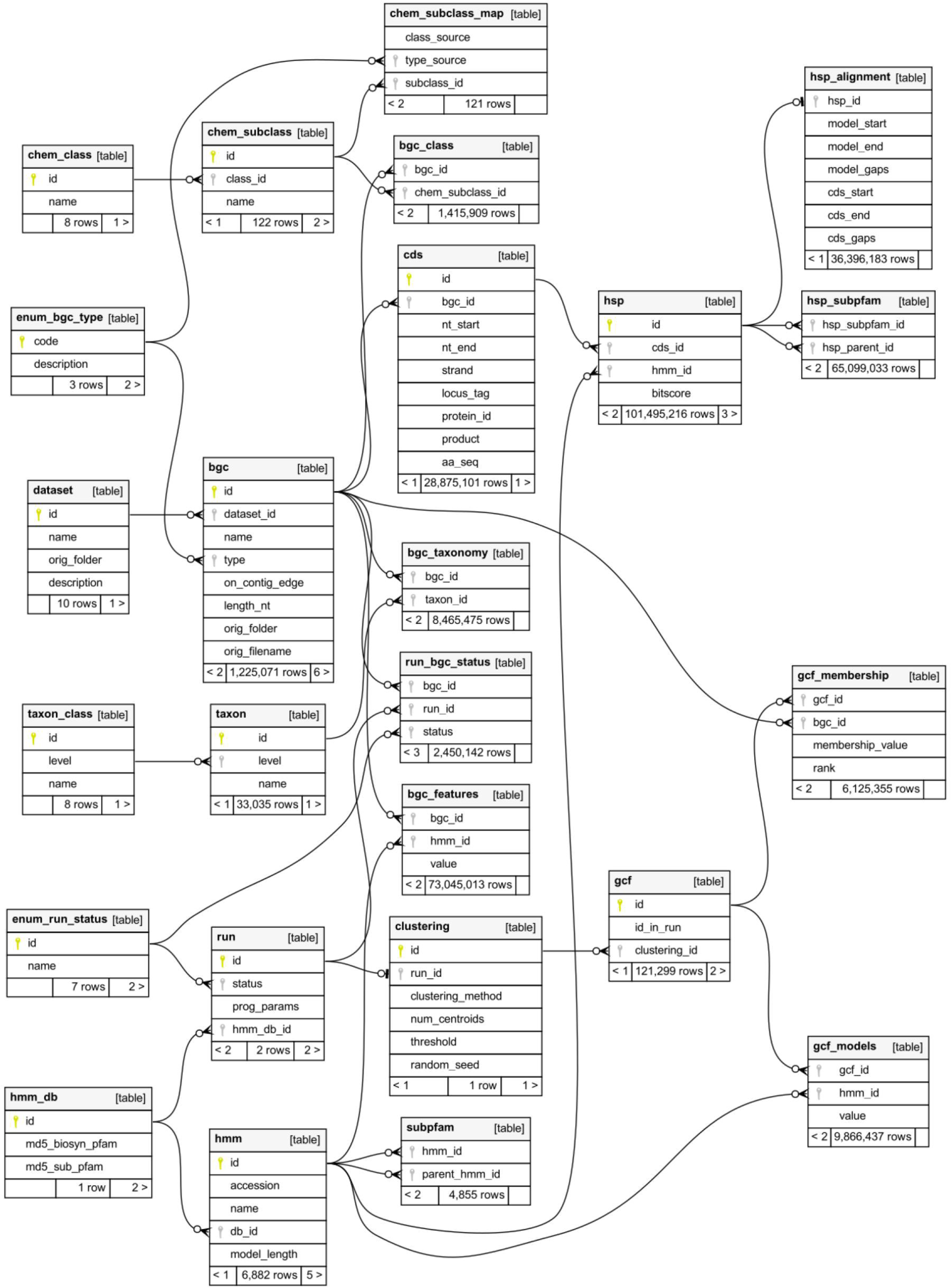
An Entity-Relationship Diagram (ERD) of the SQLite3 database used in BiG-SLiCE v1.0.0 (this study). The ERD was generated using SchemaSpy version 6.1.0 (http://schemaspy.org/).

**Supplementary Figure 3.**
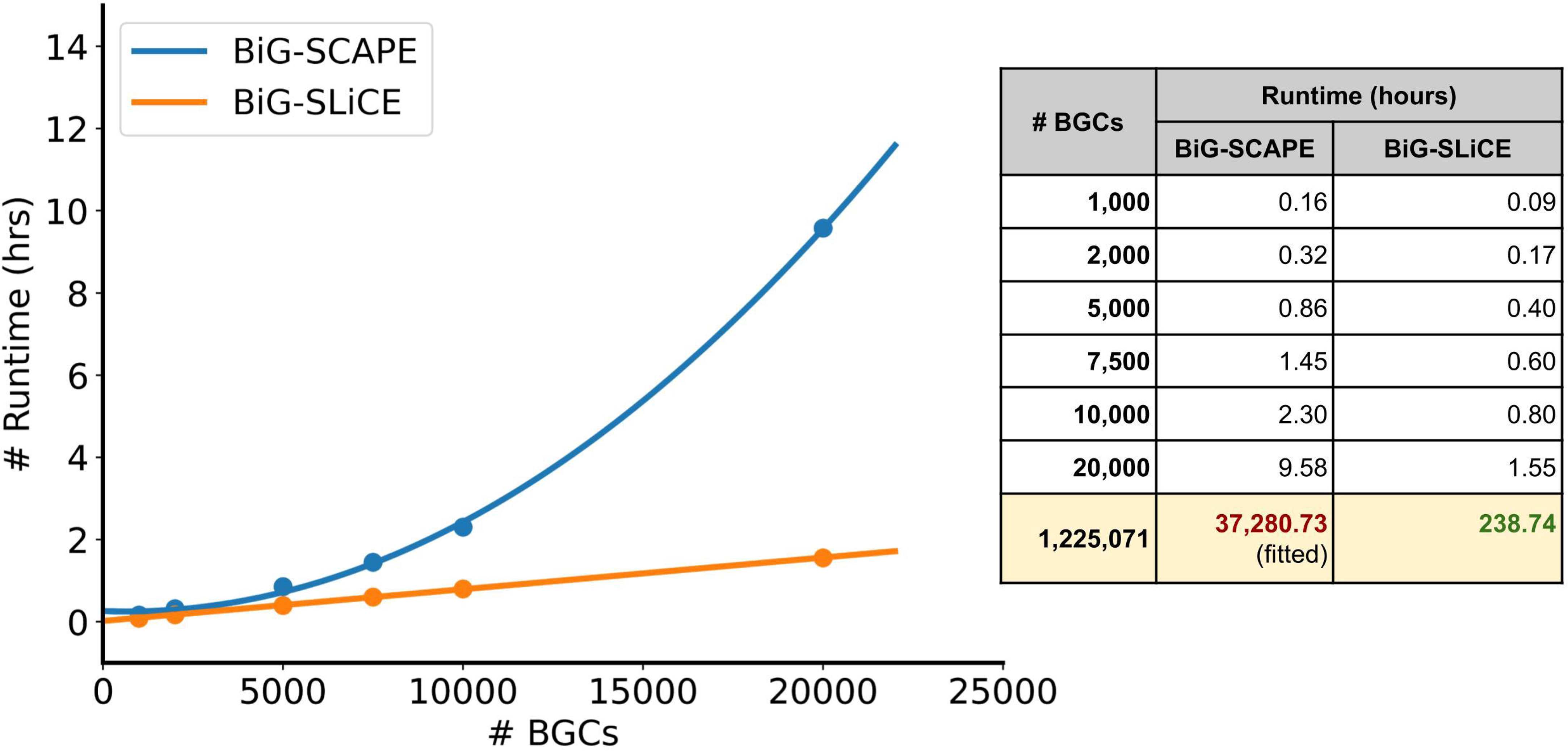
Runtime comparison between BiG-SCAPE and BiG-SLiCE. Runs were performed on a 36-cores CPU using subsets of randomly sampled BGCs from the dataset (a single subset will be used for both compared runs and will also be included for subsequent runs with larger subsets). Using data points from the sampled runs, a curve was fitted to estimate the runtime of an input size of 1,225,071 BGCs for BiG-SCAPE, while the real runtime taken from the full run log of *T* = 300 is used for BiG-SLiCE.

**Supplementary Figure 4.**
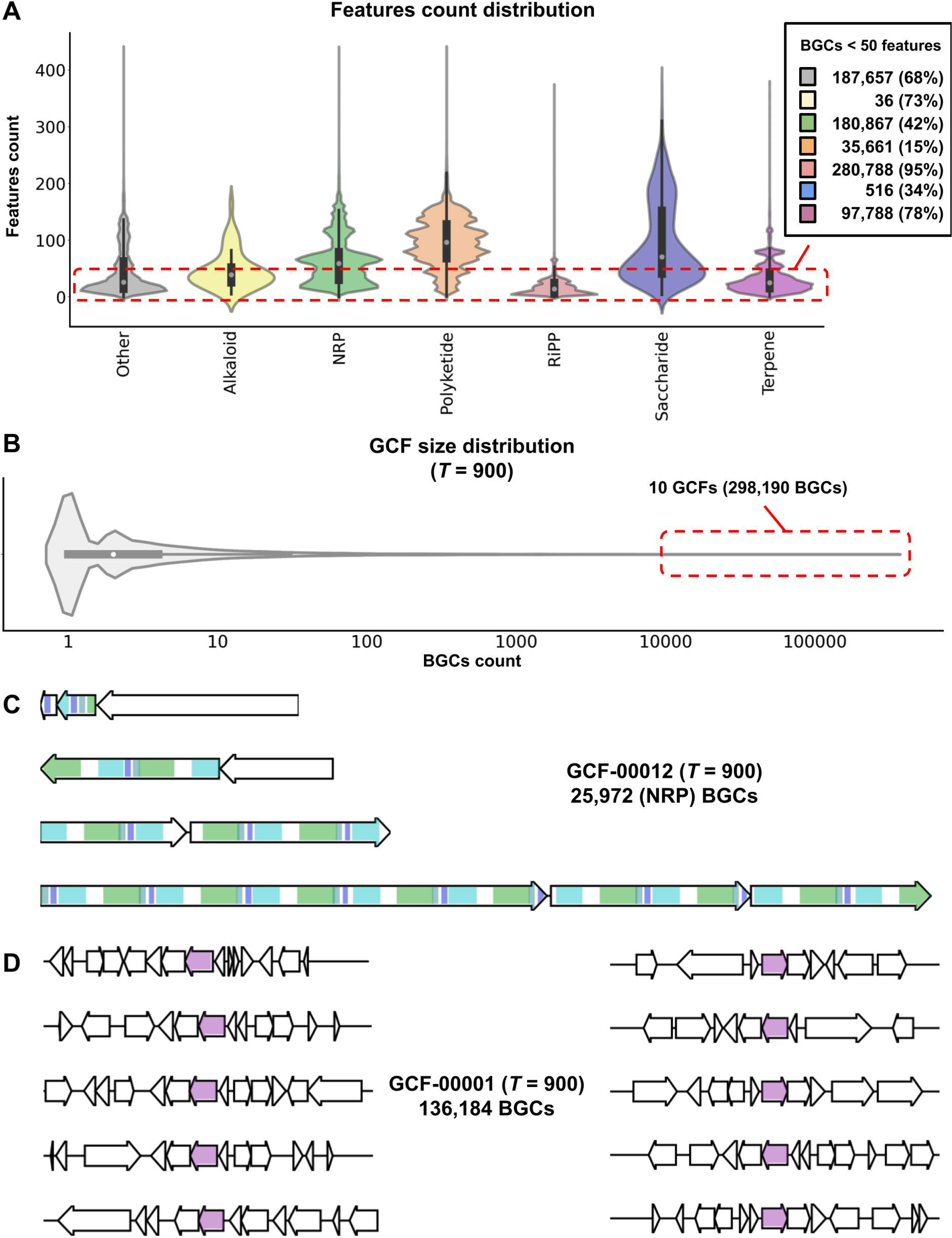
**A.** Distribution of features count (calculated by the total feature values divided by 255) across different BGC classes. Here, the distribution of BGCs having less than 50 features is highlighted, showing that some BGC classes tend to have much fewer features than others. **B.** Distribution of GCF sizes from the *T* = 900, showing some GCFs having a significantly high number of BGCs, mainly due to the effect of low features count of the BGCs. **C.** Examples of BGCs having high copy numbers of the same domain, and **D.** BGCs relying on (or having) only a single biosynthetic domain as detected by BiG-SLiCE, thus resulting in a highly similar features matrix, leading them being grouped together into a single GCF.

### Supplementary tables

**Supplementary table 1.** List of biosynthetic-Pfam pHMMs used by BiG-SLiCE.

**Supplementary table 2.** List of “core” biosynthetic-Pfam and the respective sub-Pfam pHMM models.

**Supplementary table 3.** List of genomes per dataset along with the total count of BGCs predicted by antiSMASH and their assigned taxonomy.

**Supplementary table 4.** Summary of five different run parameters on the full dataset of 1.2M BGCs.

**Supplementary table 5.** Calculated statististics of the 266 GCFs that were used to annotate the global phylogram map of biosynthetic diversity.

**Supplementary table 6.** BGC counts per dataset of 29,955 GCFs from the *T* = 900 run and the calculated distance to the closest matching MIBiG BGC.

**Supplementary table 7.** Unique GCF counts of species having at least 4 strain genomes in the full dataset.

**Supplementary table 8.** BGC class absence/presence distribution of species in the full dataset. Hybrid BGCs will have each of their classes counted separately, meaning the sum of the numbers will not be equal to the total number of BGCs per species.

## Abbreviations

AMR: antimicrobial resistant
BGC: biosynthetic gene cluster
GCF: gene cluster family
GTDB: Genome Taxonomy Database
MAG: metagenome-assembled genome
NCBI: National Center for Biotechnology Information
NRPS: non-ribosomal peptide synthase
pHMM: protein hidden markov model
PKS: polyketide synthase
RiPP: ribosomally translated post-translationally modified peptide

## Acknowledgements

We thank Joris Louwen for adding 21 manually selected biosynthetic pfams to the library, Vittorio Tracanna for his constructive feedback on the study, and Jorge C. Navarro-Muñoz for his input on BiG-SCAPE and fungal BGCs.

## Competing Interests

M.H.M. is a co-founder of Design Pharmaceuticals and a member of the scientific advisory board of Hexagon Bio.

## Funding

The work of S.A.K. was supported by the Graduate School for Experimental Plant Sciences (EPS), The Netherlands. J.J.J.v.d.H. acknowledges funding by the Netherlands eScience Center (NLeSC) Accelerating Scientific Discoveries Grant [ASDI.2017.030].

## Author’s Contributions

SAK and MHM conceived the study. SAK designed and wrote the BiG-SLiCE software. SAK collected and processed all input data. SAK performed all analyses with the help and input from all other authors. JJJVDH and MHM provided input on the biochemical perspective of the study. DDR, JJJVDH and MHM provided input on the computational parts of the clustering algorithm. SAK wrote the initial draft of the paper. All authors contributed to writing and editing the final version of the manuscript.

## Notes

https://zenodo.org/record/3975432

### Summary of Updates

fix GitHub repository URL

