## Supplementary texts for "BiG-SLiCE: A Highly Scalable Tool Maps the Diversity of 1.2 Million Biosynthetic Gene Clusters"

**Supplementary Text 1.**

NCBI query scripts (to be used in <https://www.ncbi.nlm.nih.gov/assembly/advanced/>) used to download all isolate genomes for this study:

- **Isolate_archaea:** "archaea"[Organism] AND ("latest genbank"[filter] AND (all[filter] NOT "derived from surveillance project"[filter] AND all[filter] NOT anomalous[filter] AND all[filter] NOT "untrustworthy as type"[Excluded from RefSeq] AND all[filter] NOT "missing trna genes"[filter] AND all[filter] NOT "missing rrna genes"[filter] AND all[filter] NOT "missing ribosomal protein genes"[filter] AND all[filter] NOT "many frameshifted proteins"[filter] AND all[filter] NOT "low quality sequence"[filter] AND all[filter] AND all[filter] NOT "low gene count"[filter] AND all[filter] AND all[filter] NOT "genome length too small"[filter] AND all[filter] NOT partial[filter] AND all[filter] NOT "abnormal gene to sequence ratio"[filter] AND all[filter] NOT "derived from environmental source"[filter] AND all[filter] NOT "derived from metagenome"[filter] AND all[filter] NOT "genome length too large"[filter]))
- **Isolate_bacteria_complete:** "bacteria"[Organism] AND ("latest refseq"[filter] AND ("complete genome"[filter] OR "chromosome level"[filter]) AND (all[filter] NOT "derived from surveillance project"[filter] AND all[filter] NOT anomalous[filter] AND all[filter] NOT "abnormal gene to sequence ratio"[filter] AND all[filter] NOT "derived from environmental source"[filter] AND all[filter] NOT "missing ribosomal protein genes"[filter] AND all[filter] NOT "untrustworthy as type"[Excluded from RefSeq] AND all[filter] NOT "unverified source organism"[filter] AND all[filter] NOT "sequence duplications"[filter] AND all[filter] NOT "missing rrna genes"[filter] AND all[filter] NOT "derived from single cell"[filter] AND all[filter] NOT "derived from metagenome"[filter] AND all[filter] NOT "genome length too large"[filter] AND all[filter] NOT "genome length too small"[filter] AND all[filter] AND all[filter] AND all[filter] NOT "low gene count"[filter] AND all[filter] NOT "low quality sequence"[filter] AND all[filter] NOT "many frameshifted proteins"[filter] AND all[filter] NOT "missing trna genes"[filter] AND all[filter] NOT chimeric[filter] AND all[filter] NOT contaminated[filter] AND all[filter] NOT misassembled[filter] AND all[filter] NOT "mixed culture"[filter] AND all[filter] NOT partial[filter]))
- **Isolate_bacteria_draft:** "bacteria"[Organism] AND ("latest refseq"[filter] AND ("scaffold level"[filter] OR "contig level"[filter]) AND (all[filter] NOT "derived from surveillance project"[filter] AND all[filter] NOT anomalous[filter] AND all[filter] NOT "abnormal gene to sequence ratio"[filter] AND all[filter] NOT "derived from environmental source"[filter] AND all[filter] NOT "missing ribosomal protein genes"[filter] AND all[filter] NOT "untrustworthy as type"[Excluded from RefSeq] AND all[filter] NOT "unverified source organism"[filter] AND all[filter] NOT "sequence duplications"[filter] AND all[filter] NOT "missing rrna genes"[filter] AND all[filter] NOT "derived from single cell"[filter] AND all[filter] NOT "derived from metagenome"[filter] AND all[filter] NOT "genome length too large"[filter] AND all[filter] NOT "genome length too small"[filter] AND all[filter] AND all[filter] AND all[filter] NOT "low gene count"[filter] AND all[filter] NOT "low quality sequence"[filter] AND all[filter] NOT "many frameshifted proteins"[filter] AND all[filter] NOT "missing trna genes"[filter] AND all[filter] NOT chimeric[filter] AND all[filter] NOT contaminated[filter] AND all[filter] NOT misassembled[filter] AND all[filter] NOT "mixed culture"[filter] AND all[filter] NOT partial[filter]))
- **Isolate_fungi:** "fungi"[Organism] AND ("latest genbank"[filter] AND (all[filter] NOT "derived from surveillance project"[filter] AND all[filter] NOT anomalous[filter] AND all[filter] NOT "untrustworthy as type"[Excluded from RefSeq] AND all[filter] NOT "missing trna genes"[filter] AND all[filter] NOT "missing rrna genes"[filter] AND all[filter] NOT "missing ribosomal protein genes"[filter] AND all[filter] NOT "many frameshifted proteins"[filter] AND all[filter] NOT "low quality sequence"[filter] AND all[filter] AND all[filter] NOT "low gene count"[filter] AND all[filter] AND all[filter] NOT "genome length too small"[filter] AND all[filter] NOT partial[filter] AND all[filter] NOT "abnormal gene to sequence ratio"[filter] AND all[filter] NOT "derived from environmental source"[filter] AND all[filter] NOT "derived from metagenome"[filter] AND all[filter] NOT "genome length too large"[filter]))
